# Hypoxia-Induced Modulation of Cellular and in vivo uptake of DNA nanocages – implications in therapeutics

**DOI:** 10.64898/2026.09.02.748847

**Authors:** Sanjay Kosara, Geethu Prakash, Harsh Dave, Sivaraman Dhanasekaran, Dhiraj Bhatia

## Abstract

DNA tetrahedra (DNA Td) are promising nanocarriers for drug delivery, but how hypoxia affects their cellular internalisation remains poorly understood. We synthesised and characterised Cy5-labelled DNA Td and established chemical hypoxia using 100 µM CoCl□ in HeLa, MDA-MB-231, and MCF-7 cells. Hypoxia was confirmed by HIF-1α nuclear translocation with >92% cell viability. Confocal microscopy revealed significantly reduced DNA Td uptake under hypoxia (P < 0.01), whereas transferrin and cholera toxin B uptake increased, indicating cargo-selective regulation. Temperature-arrest experiments confirmed reduced energy-dependent internalisation. Pharmacological profiling showed a shift from clathrin-mediated (42% → 29%) and galectin/glycan-dependent (43% → 35%) pathways toward lipid raft/cholesterol-dependent uptake (36% → 56%). Hypoxia increased plasma membrane electronegativity, suggesting a biophysical barrier to DNA Td uptake. Importantly, DOTMA complexation restored uptake to normoxic levels, identifying electrostatic repulsion as a key determinant. In zebrafish larvae, hypoxia significantly enhanced whole-larva DNA Td accumulation (P < 0.001). These findings highlight surface charge engineering as a strategy for improving DNA nanostructure delivery under hypoxic conditions.

## Introduction

The tumour microenvironment presents formidable barriers to therapeutic delivery. Among them, hypoxia - inadequate oxygen supply arising from aberrant tumour vasculature and rapid cellular proliferation - is a clinically significant determinant of treatment resistance and disease progression^1,2^. Hypoxia activates hypoxia-inducible factor (HIF)-driven transcriptional reprogramming that fundamentally reshapes cellular metabolism, membrane composition, and endocytic activity^3^. These adaptations profoundly alter how cells interact with incoming nanomedicines, yet the specific impact on DNA nanostructure uptake and intracellular trafficking remains largely unexplored. Addressing this gap is essential for developing nanocarriers capable of delivering therapeutic cargo within oxygen-deprived tumour regions.

DNA tetrahedra (DNA-Td) are rigid three-dimensional nanostructures self-assembled from four oligonucleotides that have emerged as versatile platforms for drug delivery and biosensing, owing to their structural stability, biocompatibility, and programmable functionality^4,5^. Unlike many synthetic nanoparticles, DNA-Td undergo efficient cellular internalisation through receptor-independent pathways, predominantly via lipid-raft-mediated endocytosis involving caveolin-1. Their compact size (typically 6–10 nm edge length), negative surface charge, and structural rigidity enable distinctive membrane interactions that facilitate cellular entry^6^. Recent studies have demonstrated their capacity to deliver therapeutic oligonucleotides, small-molecule drugs, and imaging agents to diverse cell types^7^. Critically, however, virtually all prior investigations have been conducted under normoxic conditions - leaving a fundamental knowledge gap regarding DNA-Td performance in oxygen-deprived tumour regions where therapeutic delivery is most urgently needed.

Converging evidence indicates that hypoxia substantially modulates nanoparticle–cell interactions through multiple mechanisms. Hypoxic stress alters membrane composition, particularly affecting lipid raft organisation and caveolin-1 expression^8,9^. Endocytic pathway activity becomes cargo-specific under low oxygen, with clathrin-mediated trafficking showing differential responses depending on the internalised cargo^10^. Hypoxia additionally activates endocytic recycling pathways through HIF-1-induced upregulation of Rab coupling protein (RCP), potentially reducing the net intracellular accumulation of internalised materials^11^. For DNA nanostructures specifically, preliminary observations indicate altered uptake in oxygen–glucose deprivation models^12,13^, yet systematic mechanistic characterisation of these effects - and their consequences for therapeutic delivery - remains absent.

This study addresses this critical gap by systematically investigating how hypoxic conditions influence DNA tetrahedron cellular uptake, endocytic pathway utilisation, intracellular trafficking, and subcellular localisation. We employed three cancer cell lines (HeLa, MDA-MB-231, and MCF-7), representing distinct molecular subtypes and endocytic capacities, under a CoCl-induced hypoxia model validated by HIF-1α nuclear accumulation and confirmed cell viability. Uptake was quantified by confocal laser scanning microscopy; endocytic pathway contributions were dissected using selective pharmacological inhibitors (Pitstop 2, Dynasore, MβCD, and lactose) alongside temperature-arrest assays; and cell-surface biophysics were characterised by zeta potential measurements and actin distribution analysis. Our findings reveal that hypoxia paradoxically suppresses DNA tetrahedron internalisation while increasing uptake of canonical endocytic markers - transferrin and cholera toxin B - through a reprogramming of endocytic pathway utilisation away from clathrin- and galectin/glycan-dependent routes and toward lipid raft/cholesterol-dominant uptake. This cargo-specific divergence is accompanied by hypoxia-induced increases in plasma membrane electronegativity and can be functionally reversed by DOTMA-mediated surface charge neutralisation of the DNA tetrahedron. Together, these findings establish the mechanistic basis for hypoxia-dependent modulation of DNA nanostructure uptake and identify surface charge engineering as a tractable strategy for restoring nanocarrier delivery in hypoxic tumour environments.

## Results and discussion

### Assembly, Characterization, Stability and Biocompatibility of DNA tetrahedron

DNA Tetrahedron (DNA Td) were assembled using a one-pot synthesis method^14^ from four single-stranded DNAs (ssDNA), each with 55 bases in length (**Fig. 2a)**, designed to self-assemble into a tetrahedral structure with edges of approximately 5-8 nm. Successful assembly was confirmed by electrophoretic mobility shift assay (EMSA), which showed a stepwise decrease in mobility from ssDNA (M1) to partially assembled intermediates (M1+M2 and M1+M2+M3) and the fully assembled DNA Td (M1+M2+M3+M4) (**Fig. 2b**). Atomic force microscopy was used to verify the morphology of DNA Td (**Fig. 2c-d**). Dynamic light scattering was employed to verify the hydrodynamic size of DNA Td (∼11.23±3.4 nm) (**Figure 2e**), and Zeta potential measurements showed a negative surface charge of -16.2 mV, typical for DNA-based nanostructures (**Fig. 2f**). Together, these results confirmed the successful assembly of well-defined DNA tetrahedra. To assess their suitability for cellular studies, the stability and biocompatibility of DNA Td were next evaluated. DNA Td remained stable for over 24 h at 37 °C in standard buffer and in DMEM, whereas reduced stability was observed in FBS-containing medium (**Supplementary Fig. S1d**). MTT analysis further showed no significant cytotoxicity across multiple cell lines at the tested concentrations (**Supplementary Fig. S1a-c**). Together, these results indicate that the assembled DNA Td are structurally well-defined, stable under relevant experimental conditions, and biocompatible, making them suitable for subsequent uptake studies under normoxia and hypoxia.

**Figure 1:**
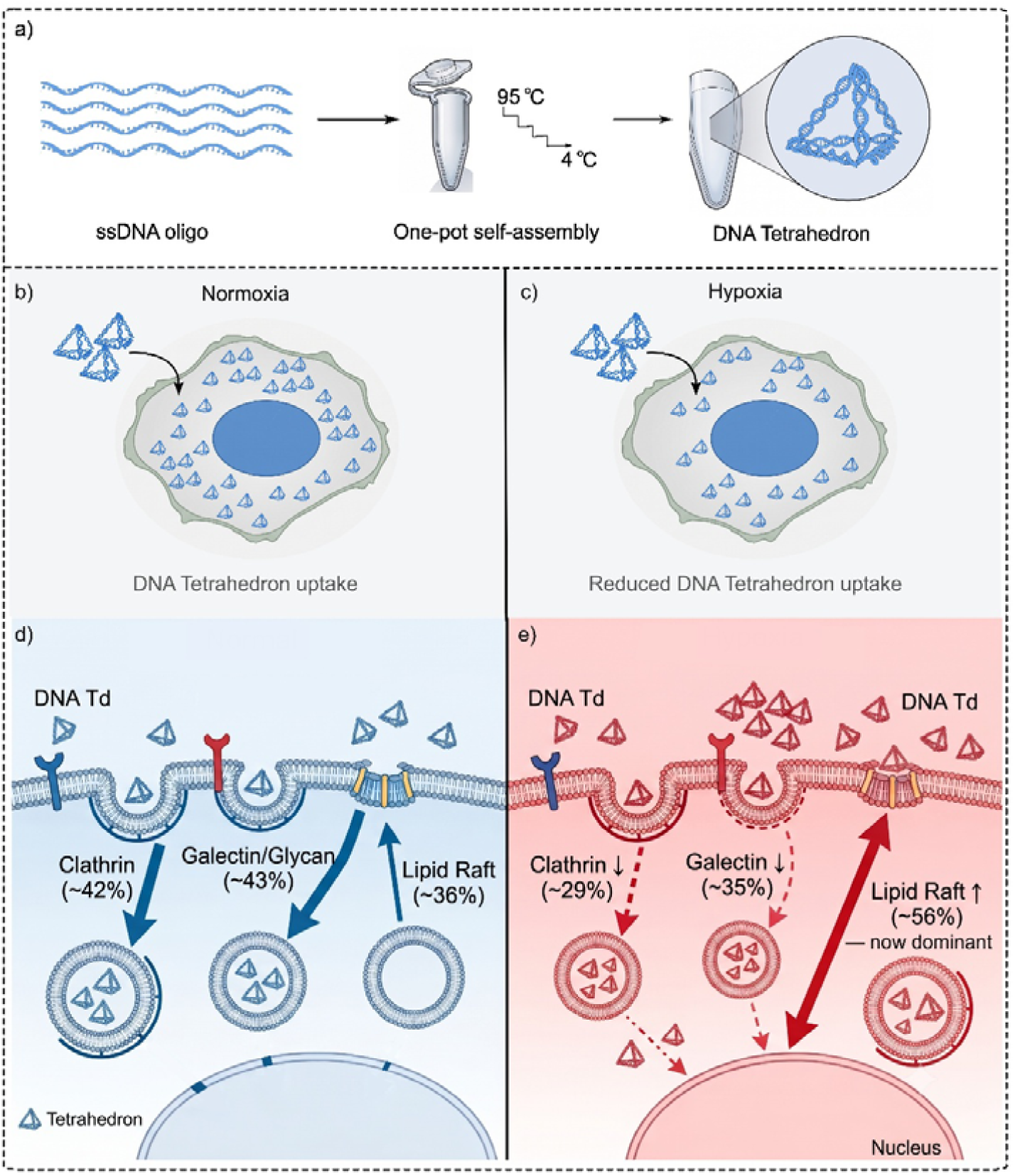
Hypoxia selectively reduces DNA tetrahedron uptake by remodelling endocytic pathways. (a) DNA tetrahedra (DNA Td) were synthesized by one-pot thermal annealing of four ssDNA oligonucleotides. (b,c) DNA Td uptake is high under normoxia but significantly reduced under hypoxia. (d,e) Under normoxia, DNA Td internalization occurs through clathrin-mediated (∼42%), galectin/glycan-dependent (∼43%), and lipid raft-mediated (∼36%) endocytosis. Under hypoxia, clathrin (∼29%) and galectin/glycan (∼35%) pathways are reduced, whereas lipid raft-mediated uptake increases (∼56%) and becomes the predominant route, despite an overall decrease in DNA Td internalisation.

**Figure 2.**
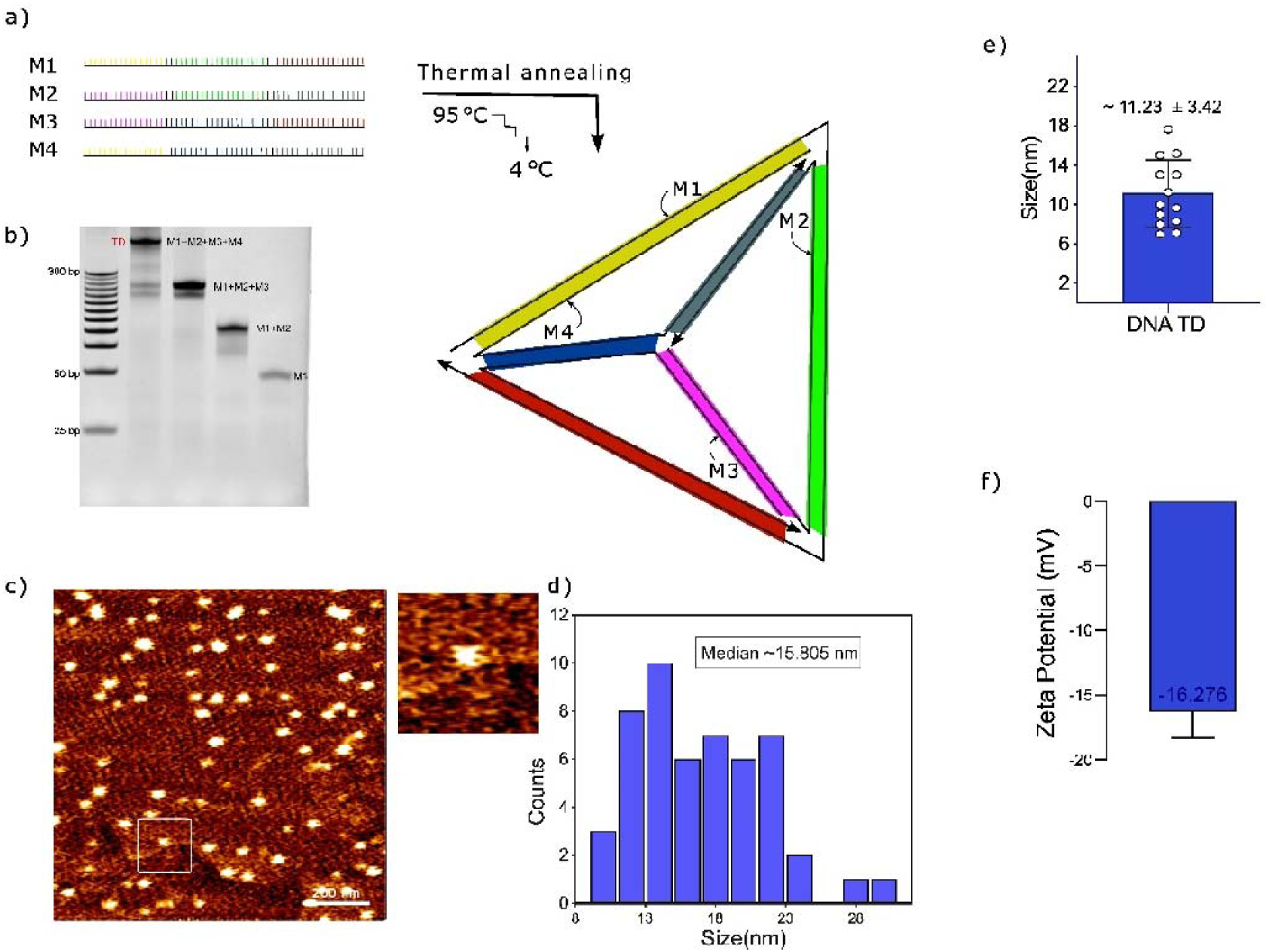
Assembly, characterization, stability, and biocompatibility of DNA tetrahedron. (a) Schematic representation of the one-pot assembly of DNA tetrahedra (DNA Td) from four single-stranded DNAs (M1-M4), each 55 nucleotides in length. (b) Electrophoretic mobility shift assay (EMSA) showing the stepwise assembly of DNA Td. A progressive decrease in electrophoretic mobility was observed from individual ssDNA (M1) to partially assembled intermediates (M1+M2, M1+M2+M3) and the fully assembled DNA Td (M1+M2+M3+M4). (c, d) Atomic force microscopy (AFM) images confirming the tetrahedral morphology of the assembled DNA Td. (e) Hydrodynamic diameter of DNA Td measured by dynamic light scattering (DLS), showing a mean size of 11.2 ± 3.4 nm. (f) Zeta potential of DNA Td, showing a surface charge of -16.3 mV. Data are presented as mean ± SD from independent experiments.

### Establishment and validation of a hypoxia model in cancer cells

To model hypoxia reproducibly, we treated three cancer cell lines (HeLa, MDA-MB-231, MCF-7), chosen for distinct molecular subtypes and endocytic behaviours, with the hypoxia mimetic CoCl□ (100 μM, 24 h), a condition previously optimised to stabilise HIF-1α while preserving viability^15^. Immunofluorescence confirmed pronounced nuclear HIF-1α accumulation under CoCl□ treatment versus low basal signal under normoxia (**Supplementary Fig. 2**), with quantification showing significantly increased nuclear HIF-1α intensity across all lines (P < 0.001; n > 50 cells/condition/line; **Supplementary Fig. 3**). All lines retained >92% viability after treatment, confirming no confounding cytotoxicity. Together, these results establish a robust, generalisable hypoxia model exhibiting HIF-1α stabilisation and nuclear localisation without loss of viability, providing a suitable framework for studying hypoxia-dependent nanomaterial uptake.

### Tumour-mimetic hypoxia differentially modulates the uptake of DNA tetrahedron and endocytic markers

Having established a validated hypoxia model, we next investigated whether tumour-mimetic hypoxia influences the cellular uptake of DNA tetrahedra. DNA Td were incubated with normoxic and hypoxic HeLa, MDA-MB-231, and MCF-7 cells, and uptake was assessed by Confocal laser scanning microscope (CLSM) and quantitative image analysis at 30 min **(Fig. 3a-c)**. Additional time points (15 and 60 min) showed consistent trends and are provided in **Supplementary Figure S4**. Under hypoxic conditions, DNA Td uptake was significantly reduced compared with normoxic controls across all three cell lines **(Fig. 3d-f)**. Quantitative analysis confirmed a marked decrease in intracellular fluorescence intensity in hypoxic cells relative to their normoxic counterparts. The reduction was consistent regardless of the distinct molecular backgrounds and endocytic capacities of the tested cell lines, suggesting that hypoxia-associated suppression of DNA Td uptake is a generalisable phenomenon. To determine whether this reduction reflected a global impairment of endocytosis, uptake of two well-characterised endocytic markers, transferrin^16,17^ and cholera toxin subunit B (CTxB)^18^, was examined under identical conditions in HeLa cells **(Fig. 3g-j)**. In contrast to DNA Td, transferrin uptake was higher under hypoxia than under normoxia. CTxB uptake showed an even more pronounced increase, consistent with enhanced caveolar or lipid raft-mediated internalisation under hypoxic conditions.

**Figure 3.**
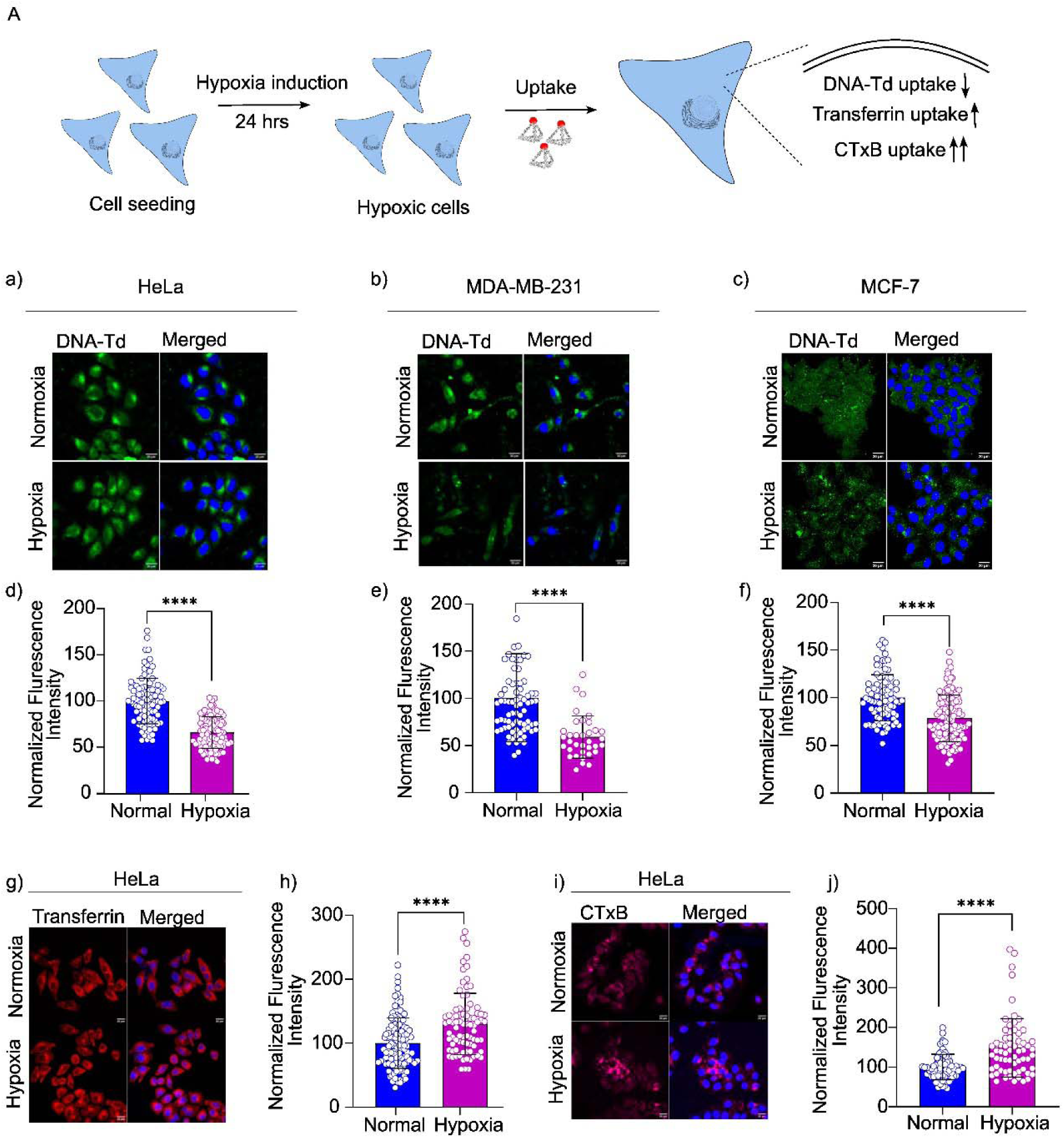
Hypoxia differentially modulates the uptake of DNA tetrahedra and endocytic markers in cancer cells. (A) Schematic representation of the experimental workflow for assessing DNA tetrahedron (DNA Td) uptake under normoxic and hypoxic conditions. (a-c) Representative CLSM images showing DNA tetrahedron (DNA Td) uptake in (a) HeLa, (b) MDA-MB-231, and (c) MCF-7 cells under normoxic and hypoxic conditions at 30 min. Scale bar: 20 μm. (d-f) Quantification of intracellular DNA Td fluorescence intensity in (d) HeLa, (e) MDA-MB-231, and (f) MCF-7 under normoxia and hypoxia. Data are presented as mean ± SD from three independent experiments. ****P < 0.001. (g, h) Representative CLSM images (g) and quantification (h) of transferrin uptake in HeLa cells under normoxia and hypoxia, showing increased uptake under hypoxic conditions. (i, j) Representative CLSM images (i) and quantification (j) of cholera toxin subunit B (CTxB) uptake in HeLa cells under normoxia and hypoxia, showing a pronounced increase under hypoxic conditions.

Together, these findings demonstrate that tumour-mimetic hypoxia does not uniformly suppress endocytosis but instead differentially modulates uptake in a cargo-dependent manner. While DNA Td internalisation is impaired, the uptake of transferrin and CTxB is enhanced, indicating that the hypoxia-associated reduction in DNA Td uptake reflects altered cellular uptake dynamics rather than a general endocytic shutdown.

### Endocytic Pathway Inhibitor Studies Reveal Pathway-Specific Modulation by Hypoxia

To probe the mechanism underlying reduced DNA Td uptake under hypoxia, we first assessed energy dependence by comparing internalisation at 37°C and 4°C - conditions that permit and block active endocytosis, respectively - under both normoxic and hypoxic conditions **(Fig. 4a, b)**. Under normoxia, DNA Td fluorescence was significantly lower at 4°C than at 37°C, confirming that internalisation proceeds via an active, energy-dependent process. Under hypoxia, this relationship was strikingly reversed: fluorescence was significantly higher at 4°C than at 37°C, indicating suppression of active internalisation and a relative predominance of passive surface binding. This reversal was cargo-specific. Transferrin maintained energy-dependent uptake under both normoxia and hypoxia (**Fig. 4c, d**), confirming that clathrin-mediated endocytosis remains operative under hypoxia. CTxB showed the expected energy-dependence under normoxia (**Fig. 4e, f**) but no significant temperature-dependent difference under hypoxia, consistent with a shift toward constitutively active lipid raft dynamics at the hypoxic membrane.

**Figure 4.**
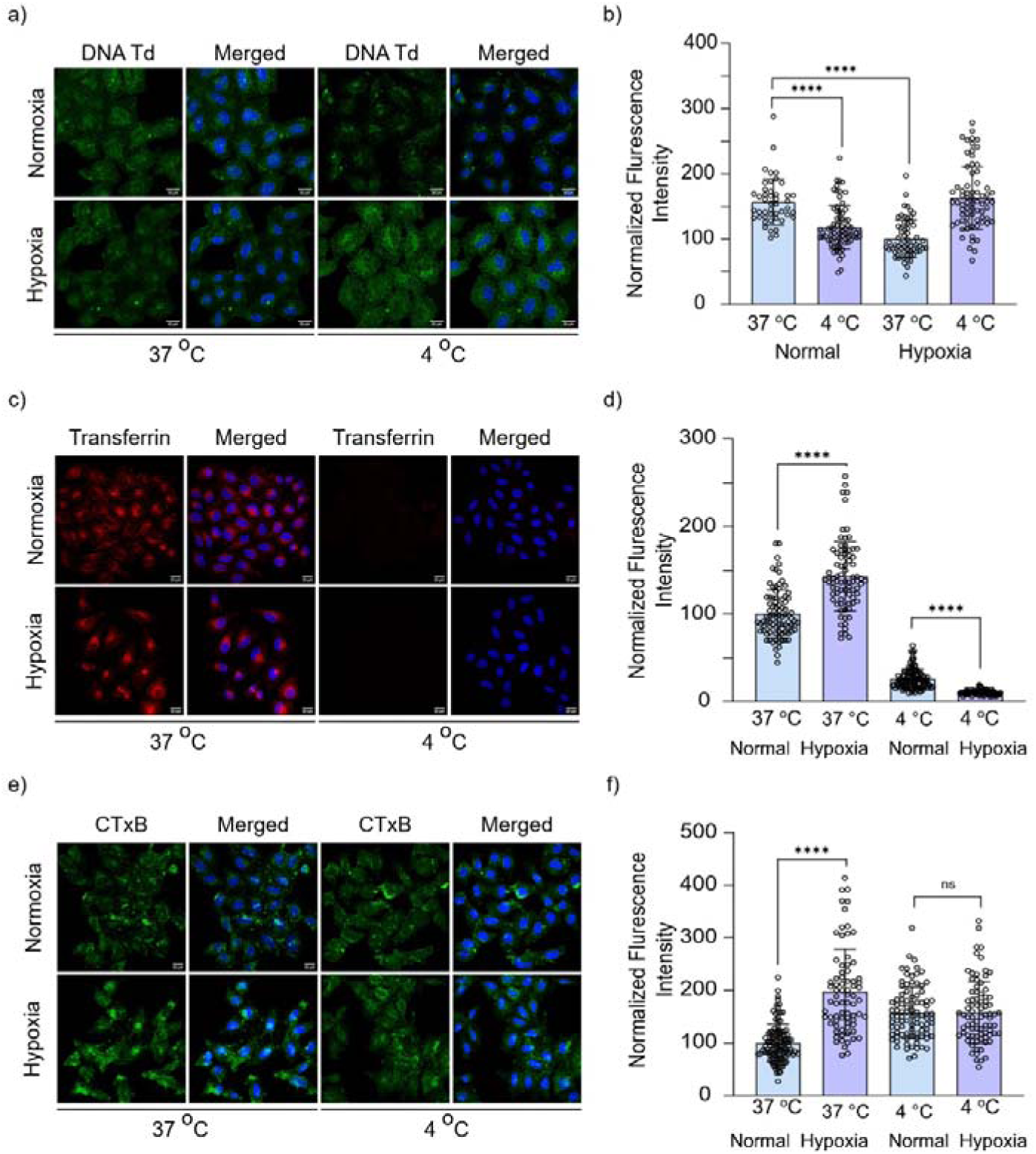
Hypoxia reverses the energy-dependence of DNA tetrahedron uptake. (a) CLSM images of DNA Td (green) and DAPI (blue) in HeLa cells at 37°C and 4°C under normoxia and hypoxia. Scale bars, 20 μm. (b) DNA Td fluorescence is lower at 4°C vs 37°C in normoxia (****P < 0.0001) but higher at 4°C in hypoxia (****P < 0.0001), showing a reversal. (c) CLSM images of transferrin (red) and DAPI (blue). Scale bars, 20 μm. (d) Transferrin is lower at 4°C vs 37°C under both conditions (****P < 0.0001), indicating energy-dependent endocytosis remains. (e) CLSM images of CTxB (green) and DAPI (blue). Scale bars, 20 μm. (f) CTxB is lower at 4°C vs 37°C in normoxia (****P < 0.0001), but no difference under hypoxia. Data are mean ± SD; n ≥ 3, ≥50 cells per condition. Two-way ANOVA with Šídák’s test. ****P < 0.0001; ns

To identify the specific pathways involved, we applied four selective pharmacological inhibitors - Pitstop 2 (clathrin-mediated endocytosis)^19^, Dynasore (dynamin-dependent fission)^6,20^, MβCD (lipid raft/cholesterol-dependent endocytosis)^21,22^, and lactose (galectin/glycan-mediated endocytosis)^23,24^ and quantified DNA Td uptake by CLSM under normoxic and hypoxic conditions (**Fig. 5a, b**). Under hypoxia, the inhibitory effect of Pitstop 2 was significantly reduced relative to normoxia (−12.98%, **Fig. 5c**), as was that of lactose (−8.29%, **Fig. 5f**), indicating diminished contributions from clathrin-mediated and galectin/glycan-dependent routes. Dynasore produced comparable inhibition under both conditions (**Fig. 5d**), By contrast, MβCD-mediated inhibition was significantly greater under hypoxia (+19.73%, **Fig. 5e**), indicating that cholesterol-dependent lipid raft endocytosis is the dominant internalisation route under low-oxygen conditions. Two-way ANOVA confirmed a significant oxygen-condition × inhibitor interaction (F(3,540) = 29.95, P < 0.0001) and a significant inhibitor main effect (F(3,540) = 32.26, P < 0.0001); the absence of a significant oxygen main effect (F(1,540) = 0.43, P = 0.51) indicates that hypoxia exerts pathway-differential rather than uniform effects on total uptake (**Fig. 5g**). Collectively, these data establish that hypoxia reprograms DNA Td internalisation from clathrin/galectin-dominant routes toward lipid raft/cholesterol-dependent uptake (**Fig. 5h**) a pathway shift that mechanistically explains the cargo-divergent responses first observed in Figure 3 and the enhanced CTxB uptake seen under hypoxic conditions.

**Figure 5.**
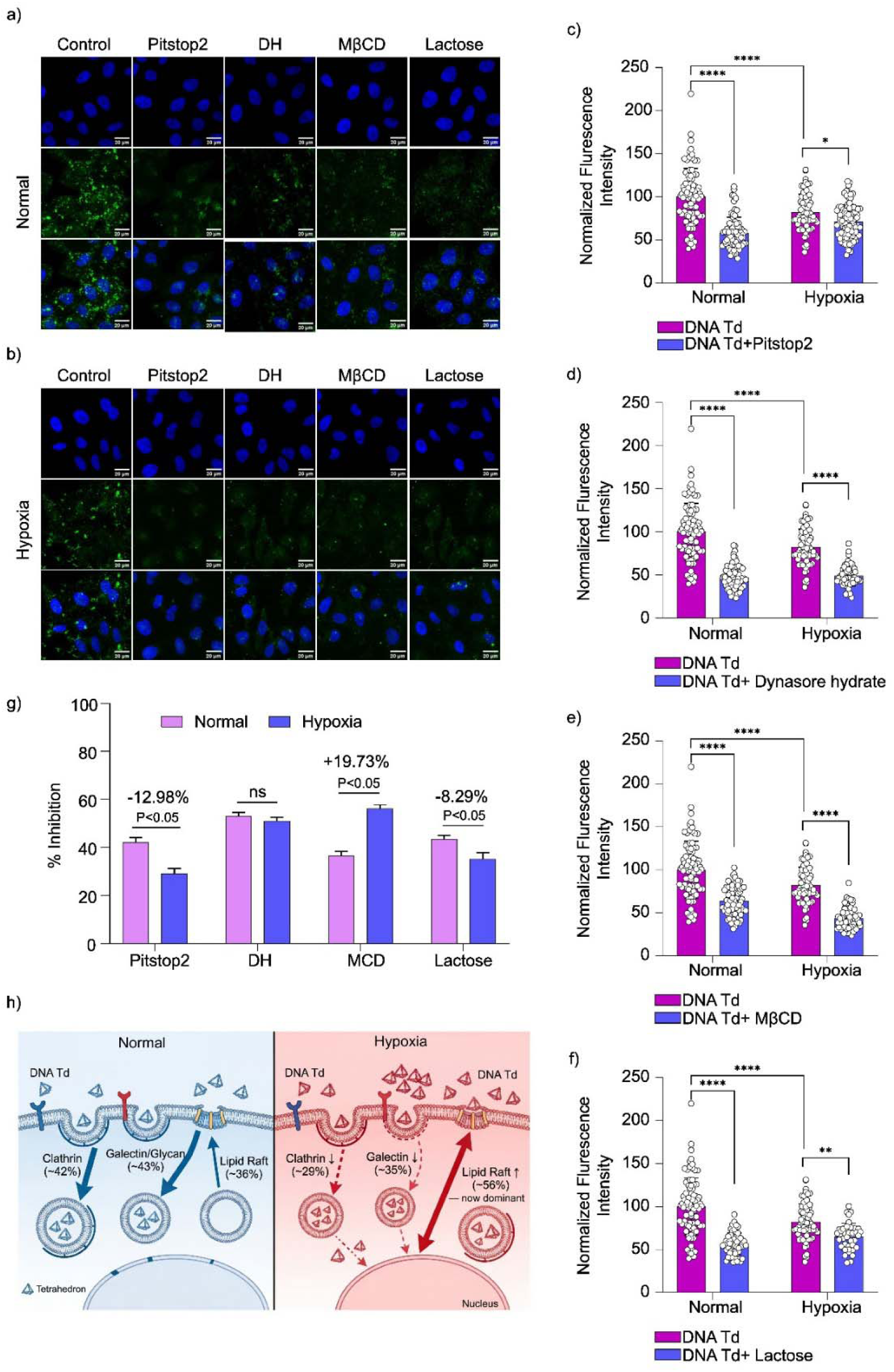
Hypoxia reprograms DNA tetrahedron internalization from clathrin/galectin-dominant to lipid raft-dependent pathways. (a,b) Representative CLSM images (DAPI, DNA Td, merged) of HeLa cells under normoxia (a) and hypoxia (b) following treatment with vehicle control, Pitstop2, Dynasore hydrate (DH), MβCD, or lactose. Scale bars, 20 μm. (c) Pitstop2 inhibition significantly reduced under hypoxia vs normoxia (−12.98%, *P < 0.05). (d) Dynasore hydrate inhibition similar under both conditions (ns). (e) MβCD inhibition significantly greater under hypoxia (+19.73%, ****P < 0.0001). (f) Lactose inhibition significantly reduced under hypoxia (−8.29%, ****P < 0.0001). (g) Summary % inhibition bar graph. Two-way ANOVA: interaction F (3,540) = 29.95, P < 0.0001; inhibitor F(3,540) = 32.26, P < 0.0001; oxygen F(1,540) = 0.43, P = 0.5129. Post hoc Šídák: Pitstop2 −12.98% (P < 0.05), Dynasore ns, MβCD +19.73% (P < 0.05), lactose −8.29% (P < 0.05). (h) Mechanistic schematic: normoxia (blue) clathrin (∼42%), galectin/glycan (∼43%), lipid raft (∼36%); hypoxia (red) clathrin (↓∼29%), galectin/glycan (↓∼35%), lipid raft dominant (↑∼56%). Data are mean ± SEM; n ≥ 3 independent experiments, ≥50 cells per condition. Magenta = DNA Td alone; blue = DNA Td + inhibitor. ****P < 0.0001; **P < 0.01; *P < 0.05; ns, not significant.

### Surface charge modulation partially restores DNA tetrahedron uptake under hypoxia

Having established that hypoxia reprograms DNA Td internalisation toward lipid raft-dependent pathways (Fig. 5), we investigated whether changes in cell-surface biophysics contribute to this shift, focusing on plasma membrane surface charge. Zeta potential measurements revealed that hypoxia increased membrane electronegativity from −10.58 mV to −12 mV (**Supplementary Figure 5**). Given the substantial negative surface charge of DNA Td (−16.2 mV; Fig. 2), this shift is expected to strengthen electrostatic repulsion at the Td–membrane interface and raise the barrier to membrane contact.

To directly test the functional role of surface charge, DNA Td was complexed with the cationic lipid DOTMA^25,26^. Gel retardation confirmed efficient complexation at 200 nM DOTMA (**Fig. 6b**), and zeta potential measurements across TD:DOTMA ratios showed a progressive shift from ∼−20 mV (1:20) to ∼+15 mV (1:250), with charge neutralisation near 1:200 (**Fig. 6c**). The 1:100 ratio was selected for cellular uptake experiments. CLSM showed markedly higher intracellular fluorescence for DNA Td–DOTMA under both conditions (**Fig. 6d**), with quantification confirming a 2–3-fold increase in hypoxic uptake and full restoration to normoxic levels (**Fig. 6e**). Together, these data demonstrate that hypoxia imposes a biophysical barrier to DNA Td internalisation through increased membrane electronegativity. The complete rescue of uptake via DOTMA complexation identifies surface charge as a tractable design parameter to improve DNA nanostructure delivery in hypoxic tumour environments.

**Figure 6.**
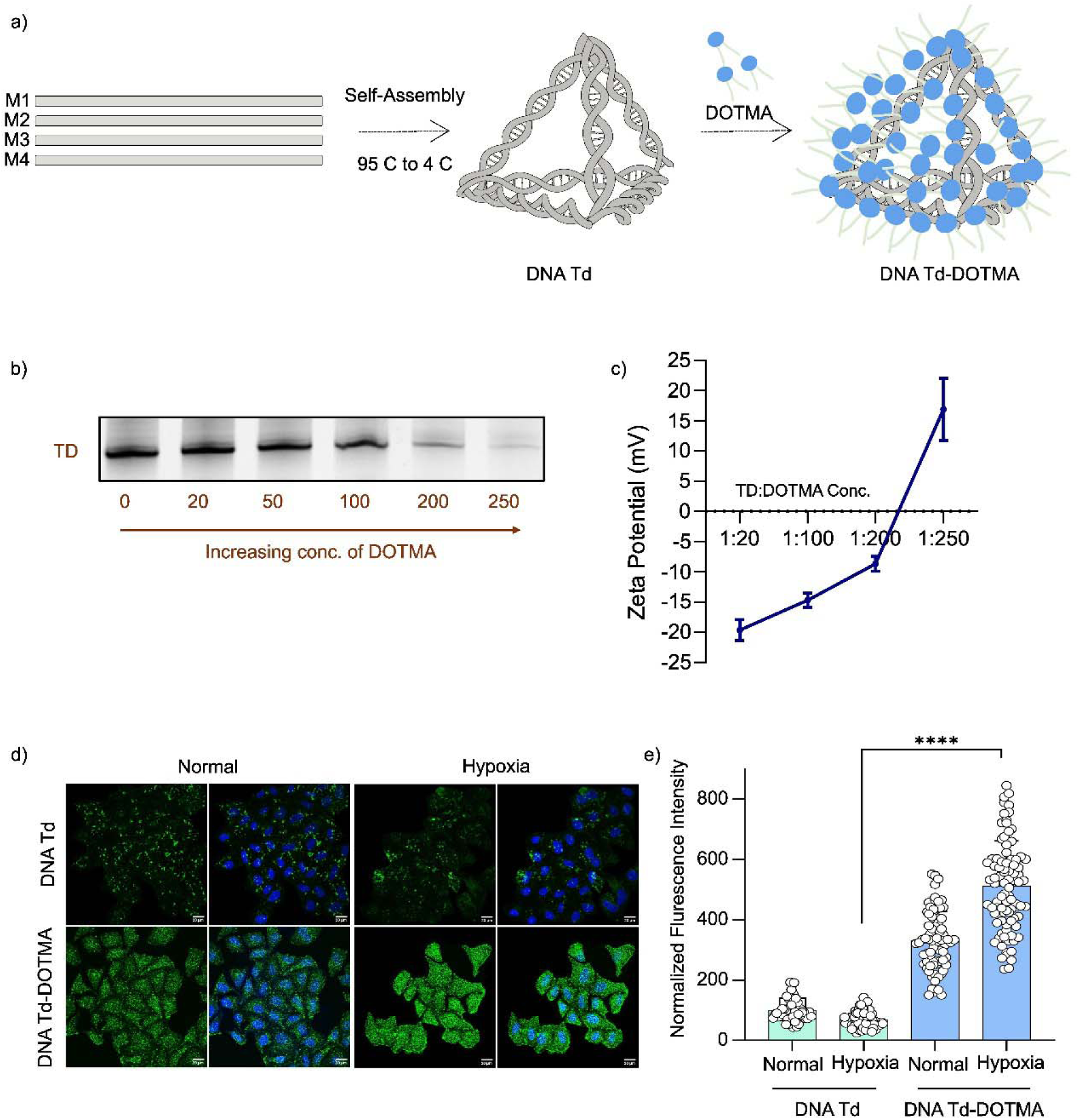
DOTMA-mediated surface charge modulation of DNA tetrahedra restores cellular uptake under hypoxia. (a) Schematic of DNA Td assembly from four oligonucleotides (M1–M4) via thermal annealing (95°C to 4°C), then DOTMA complexation to form DNA Td-DOTMA. DOTMA (blue spheres) coats the tetrahedron scaffold. (b) Agarose gel retardation assay showing the DNA Td band (labelled “TD”) at increasing DOTMA concentrations (0, 20, 50, 100, 200, 250 nM). Progressive band retardation and smearing with increasing DOTMA concentration confirms electrostatic complexation and charge neutralisation. (c) Zeta potential of DNA Td-DOTMA complexes as a function of TD:DOTMA molar ratio (1:20 to 1:250). Zeta potential shifts progressively from approximately −20 mV (at 1:20) through charge neutralisation (∼0 mV) at approximately 1:200 ratio to approximately +15 mV (at 1:250). The 1:100 ratio was selected for subsequent cellular uptake experiments. (d) Representative CLSM images of HeLa cells treated with DNA Td (top row) or DNA Td-DOTMA (bottom row) under normoxia and hypoxia. Green = DNA Td or DNA Td-DOTMA; blue = DAPI (nuclei). Scale bar, 20 μm. (e) Quantification of intracellular fluorescence intensity for DNA Td and DNA Td-DOTMA under normoxia and hypoxia. Data are mean ± SEM; two-way ANOVA with Tukey’s post-hoc test; ****P < 0.0001.

### Hypoxic Conditions Promote DNA Tetrahedron Accumulation in Zebrafish Larvae

To extend these findings to an in vivo context, 72 hpf zebrafish larvae were exposed to Cy5-labelled DNA Td under normoxic or CoCl□-induced hypoxic conditions (**Fig. 7A**). Three groups were compared: untreated controls (no DNA Td), a normoxia group exposed to DNA Td for 4 h, and a hypoxia group pretreated with CoCl□ for 1 h prior to 4 h DNA Td exposure. Whole-larva fluorescence imaging revealed minimal Cy5 signal in controls, a detectable signal in the normoxia group, and markedly stronger fluorescence throughout the hypoxia group (**Fig. 7B**). Quantification confirmed that DNA Td-associated fluorescence was significantly elevated in CoCl□-treated larvae relative to both the normoxia group and untreated controls (**Fig. 7C**), demonstrating that CoCl□-induced hypoxia substantially enhances whole-larva DNA Td accumulation in vivo.

**Figure 7.**
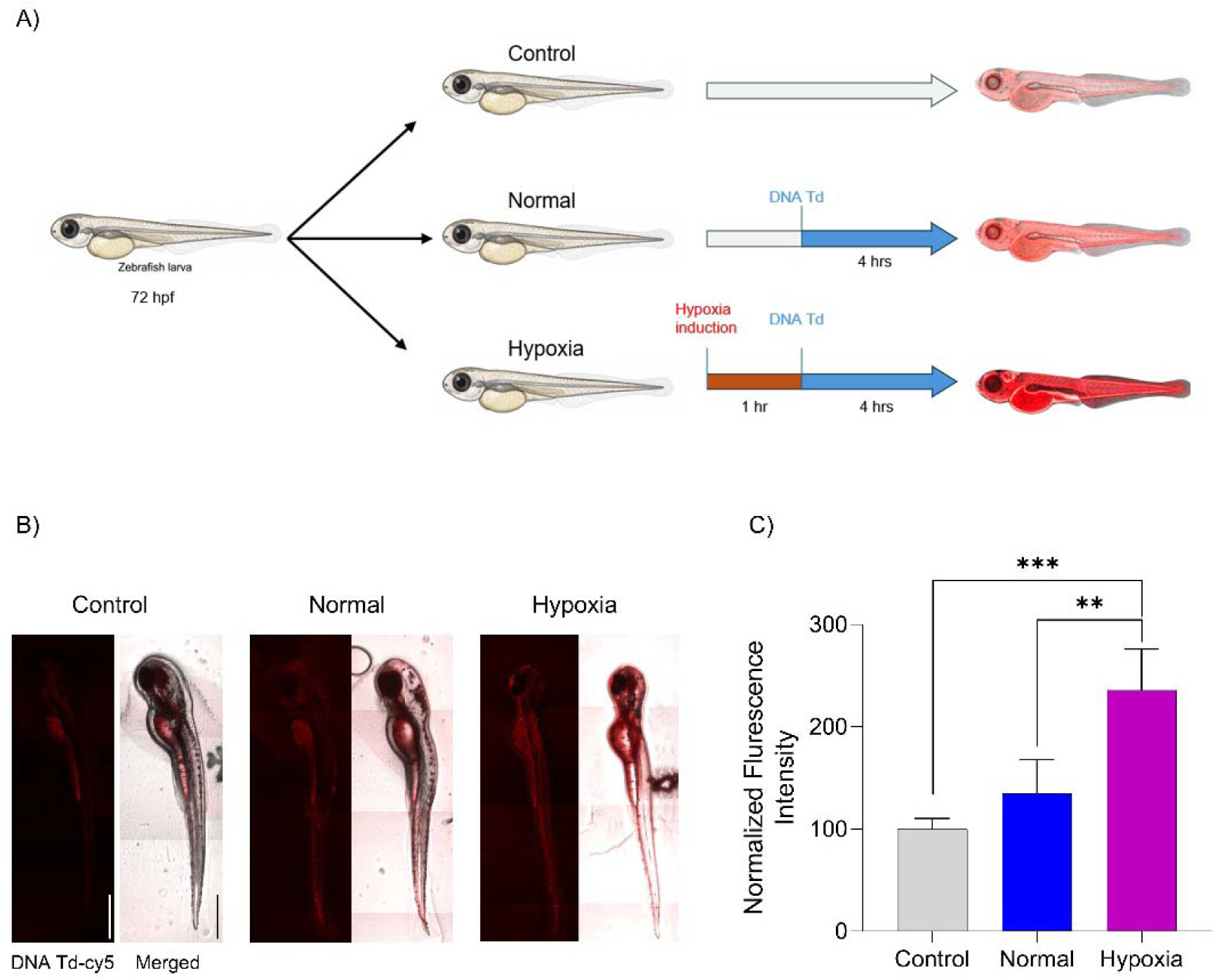
DNA tetrahedron uptake in zebrafish larvae under normoxic and hypoxic conditions. (A) Schematic representation of the experimental design using 72 hpf zebrafish larvae. Larvae were divided into control, normoxia, and CoCl -induced hypoxia groups. Hypoxia was induced by CoCl pretreatment for 1 h, followed by exposure to Cy5-labeled DNA tetrahedron (DNA-Td) for 4 h. (B) Representative fluorescence (DNA-Td-Cy5), bright-field, and merged images of zebrafish larvae showing DNA-Td distribution under control, normoxic, and hypoxic conditions. (C) Quantification of DNA-Td-associated fluorescence intensity demonstrating increased DNA-Td accumulation in hypoxic larvae compared with control and normoxic conditions. Data are presented as mean ± SD. Statistical significance was determined using one-way ANOVA with appropriate multiple-comparison testing. **p < 0.01, ***p < 0.001.

The enhanced accumulation under CoCl treatment is consistent with HIF-1α stabilisation driving coordinated changes in endocytic activity, membrane dynamics, and tissue permeability, collectively increasing DNA Td accessibility and retention within larval tissues, paralleling the pathway-level and biophysical remodelling observed in HeLa cells. However, since these measurements reflect whole-larva fluorescence, they cannot distinguish intracellular uptake from extracellular, vascular, or interstitial accumulation; the findings are therefore interpreted as enhanced DNA Td accumulation rather than definitively increased cellular internalisation. Nonetheless, these data provide initial in vivo evidence that a hypoxic microenvironment can substantially alter the biodistribution of DNA nanostructures, reinforcing the relevance of hypoxia as a microenvironmental variable in the design of nucleic acid nanocarriers for tumour-targeted delivery.

## Conclusion

This study establishes that hypoxia modulates DNA Td internalisation in a cargo-specific manner, reducing uptake in HeLa, MDA-MB-231, and MCF-7 cells even as canonical endocytic markers (transferrin, CTxB) show increased uptake under identical conditions. Mechanistically, hypoxia reprograms DNA Td internalisation from clathrin- and galectin/glycan-dependent routes toward lipid raft/cholesterol-dependent uptake, accompanied by increased plasma membrane electronegativity. DOTMA-mediated charge neutralisation fully restored DNA Td uptake to normoxic levels, identifying electrostatic repulsion at the hypoxic membrane as a primary, modifiable determinant of the uptake deficit. In vivo, CoCl -treated zebrafish larvae showed significantly enhanced whole-larva DNA Td accumulation relative to normoxic controls, apparently contrasting with the cellular uptake findings. This likely reflects HIF-1α-driven increases in tissue permeability and vascular remodelling that elevate net nanostructure retention despite per-cell internalisation deficits - a distinction that tissue-resolved imaging will be needed to resolve. Together, these findings identify surface charge engineering as a tractable strategy for restoring DNA nanostructure delivery in hypoxic tumour environments and position endocytic pathway preference as a hypoxia-tunable design parameter for nucleic acid nanocarriers. Future work should extend the pathway analysis to MDA-MB-231 and MCF-7 cells, confirm DOTMA compatibility with DNA Td cargo function, and validate these findings under physiological rather than chemically induced hypoxia.

## Methods

### Materials

Dulbecco’s Modified Eagle’s Medium (DMEM) (D5796), Fetal Bovine Serum (10270106), and trypsin-EDTA (0.25%) (15050065) are obtained from Gibco. Powder-formed ssDNA with the sequence shown in Table 1, 6X loading dye, 25 bp ladder, mowiol, transferrin-A647, CTxB, Pitstop-2, Dynasore, methyl-β-cyclodextrin and Lactose were obtained from Sigma Aldrich. Nuclease-free water and magnesium chloride are obtained from SRL, India. Acrylamide: Bisacrylamide (30 %), N,N,N′,N′-Tetramethylethylenediamine (TEMED), ammounium persulfate (APS), Penstrap, paraformaldehyde, ethidium bromide, triton X is obtained from Himedia. MTT was purchased from TCI.

**Table 1:**
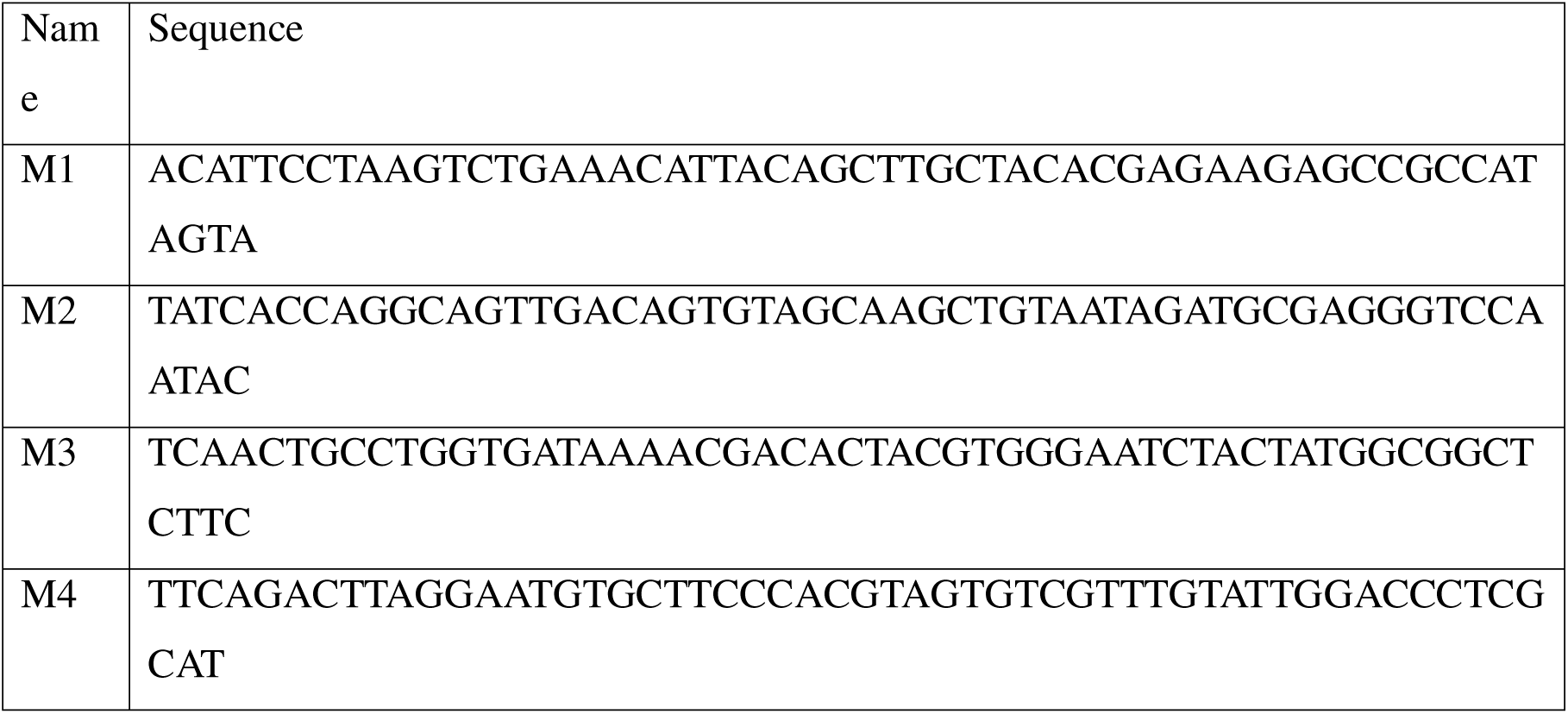
Specific DNA sequence of each single stranded DNA.

### Synthesis of DNA Tetrahedron

One-pot synthesis was used for synthesis DNA TD. In brief, the four oligonucleotides were combined with 2 mM MgCl2 in equimolar ratios (M1:M2:M3:M4). After being heated to 95°C, the reaction was gradually annealed and cooled to 4°C. The reaction cycle consisted of a 5 °C step decrease separated by 15 minutes. The final TD concentration was 2.5 μM.

### Electrophoretic Mobility. Shift Assay (EMSA)

EMSA was performed using 10% Native-PAGE at 80 V in TAE buffer for 90 min. The sample contained 5 μL of TD, 3 μL of 1×TAE loading buffer, and 1.5 μL of 6x loading dye. After electrophoresis, the bands of DNA were stained with EtBr Stain, and the image was visualized using Gel Documentation system (Biorad ChemiDoc MP Imaging System).

### Dynamic Light Scattering (DLS)

The hydrodynamic radius was measured using the Malvern Zetasizer Nano ZS. The sample was diluted to 1:20. 50 μL of a 2.5 μM DNA TD sample was used. Thirteen readings were obtained in a single run, and the measurements were made in duplicate.

### Atomic Force Microscopy (AFM)

A freshly cleaved mica was used. 5-10 μL of TD was overlaid on top of the mica sheet and allowed to dry in a desiccator, before AFM. Imaging was done using BIO-AFM (Bruker JPK NanoWizard sense+) in tapping mode, and the image was processed using JPK software.

### Cell Culture and Hypoxia Induction

HeLa (human cervical adenocarcinoma), MDA-MB-231 (human breast adenocarcinoma), and MCF-7 (human breast adenocarcinoma) cells were cultured in Dulbecco’s Modified Eagle Medium (DMEM) supplemented with 10% fetal bovine serum (FBS) and 1% penicillin-streptomycin at 37°C in a humidified atmosphere containing 5% CO□. Cells were maintained under standard culture conditions and subcultured regularly before experimental use.

For hypoxia experiments, chemical hypoxia was induced by treating cells with 100 μM cobalt(II) chloride (CoCl□, Sigma-Aldrich) for 24 h. CoCl stabilizes hypoxia-inducible factor-1α (HIF-1α) under normoxic conditions and was used as a chemical hypoxia mimic. Normoxic control cells were maintained under identical culture conditions without CoCl□. HeLa, MDA-MB-231, and MCF-7 cells were used for DNA tetrahedron uptake studies, while individual cell lines were selected for specific downstream experiments as indicated in the respective sections.

### Cellular Uptake Assays

Cells were seeded in 24-well plate at a density of 2 × 10 cells per well and allowed to adhere overnight. Cells were then subjected to normoxic or hypoxic conditions for the indicated times. For uptake experiments, cells were washed with phosphate-buffered saline (PBS) and incubated with fluorescent DNA Td-cy3/cy5 (200 nM), Tf-A488/Tf-A633 (5 μg/mL), or CtxB-A488 (1 μg/mL) in serum-free DMEM for different time points (15 mins, 30 mins, and 60 mins) at 37°C. Following incubation, cells were washed three times with 1X PBS, fixed with 4% paraformaldehyde for 15 minutes at room temperature, and washed again with PBS.

### Inhibition Studies

To investigate the endocytic pathways involved in DNA tetrahedron (DNA Td) internalisation, pharmacological inhibitors targeting distinct endocytic mechanisms were used. Cells were seeded onto glass coverslips and allowed to adhere overnight before treatment. For hypoxia experiments, cells were treated with 100 μM CoCl□ for 24 h, whereas normoxic control cells were maintained under identical conditions without CoCl□.

Prior to DNA Td treatment, cells were pre-incubated with specific inhibitors at beforehand determined concentrations. Pitstop 2 (12 μM) inhibited clathrin-mediated endocytosis; dynasore hydrate(60 μM), dynamin-dependent vesicle formation; methyl-β-cyclodextrin (MβCD)(3 mM), cholesterol-dependent lipid raft-mediated endocytosis; and lactose(80 mM), galectin/glycan-dependent uptake. After pretreatment (30 min), fluorescent DNA Td was added and incubated under the respective conditions. Control cells received DNA Td without inhibitors. Post-incubation, cells were washed with PBS, fixed with 4% paraformaldehyde, and imaged by confocal microscopy using identical settings. Confocal images were analysed in ImageJ/Fiji to quantify intracellular fluorescence after background subtraction. Inhibitor effects were calculated relative to untreated controls under normoxic or hypoxic conditions. All experiments used independent biological replicates.

### In Vivo Zebrafish Experiments

#### Danio rerio (Zebrafish) husbandry and maintenance

Adult wild-type zebrafish (Danio rerio) were maintained under standard laboratory conditions in a recirculating zebrafish housing system at 28 ± 0.5°C with a 14:10 h light cycle. Embryos were obtained through natural pairwise mating and collected within 1 h post-fertilisation (hpf). Embryos were maintained in E3 embryo medium at 28–28.5°C and staged according to standard developmental criteria. At 72 hpf, larvae were used for hypoxia induction and DNA tetrahedron (DNA-Td) uptake experiments.

#### Chemical induction of hypoxia

Chemical hypoxia was induced in 72 hpf zebrafish larvae using cobalt chloride (CoCl□). Larvae were transferred to E3 embryo medium containing the indicated concentrations of CoCl□ and incubated for 1 h at 28°C. Larvae maintained in E3 medium without CoCl□ served as the normoxic control. Following hypoxia induction, larvae underwent DNA-Td uptake analysis. The CoCl□ concentrations used to induce hypoxia were 250 µM, 500 µM, 1 mM, 5 mM, and 10 mM.

#### DNA tetrahedron uptake assay

DNA-Td uptake was evaluated using Cy5-labelled DNA tetrahedra. Following hypoxia induction, larvae were incubated with Cy5-labelled DNA-Td prepared in E3 medium for 4 h at 28°C. For the normoxic group, larvae were directly exposed to DNA-Td for 4 h without CoCl□ pretreatment. A control group without DNA-Td exposure was maintained under the corresponding experimental conditions to determine background fluorescence. Following incubation, larvae were washed thoroughly with E3 medium to remove unbound DNA-Td and subsequently processed for fluorescence imaging.

### Confocal Microscopy and Image Analysis

Confocal fluorescence microscopy was performed using a Confocal laser-scanning microscope equipped with 405 nm, 488 nm, 561 nm, and 633 nm laser lines and a 63×/1.4 NA oil-immersion objective. For each sample, z-stack images were acquired spanning the entire cell height. Imaging parameters were kept constant across all experimental conditions within each experiment to enable quantitative comparison.

Image analysis was performed using Fiji (ImageJ) software. For quantification of cellular uptake, regions of interest (ROI) were manually drawn around individual cells based on brightfield or DAPI-stained nuclear boundaries, and mean fluorescence intensity was measured for each channel. Background fluorescence was measured from cell-free regions and subtracted. For each condition, at least 50 cells from three independent experiments were analysed.

### Statistical Analysis

All experiments were performed in at least three independent biological replicates unless otherwise stated. Data are presented as mean ± standard deviation (SD) as indicated. Statistical significance was assessed using unpaired two-tailed Student’s t-tests for comparisons between two groups, or one-way analysis of variance (ANOVA) followed by Tukey’s post-hoc test for multiple comparisons. P-values < 0.05 were considered statistically significant (*P < 0.05, \**P < 0.01, P < 0.001).* All statistical analyses were performed using GraphPad Prism 9 software.

## Contributions

S.K.: Conceptualisation, Methodology, Investigation, Formal analysis, Visualisation, and Writing-original draft. G.P. and H.D.: Methodology and Investigation (zebrafish experiments). S.D.: Methodology and Resources (zebrafish experiments and zebrafish facility). D.B.: Conceptualisation, Supervision, Project administration, Writing-review & editing, and Funding acquisition. All authors reviewed and approved the final manuscript.

## Acknowledgement

All authors thank IITGN and CIF-IITGN for facilities. S.K. and G.P. thank IITGN for the fellowship. D.B. thanks SERB-DST GoI for the research grant, GUJCOST, GSBTM, and MoES-STARS for the research funding.

