## Supporting information for "Hypoxia-Induced Modulation of Cellular and in vivo uptake of DNA nanocages – implications in therapeutics"

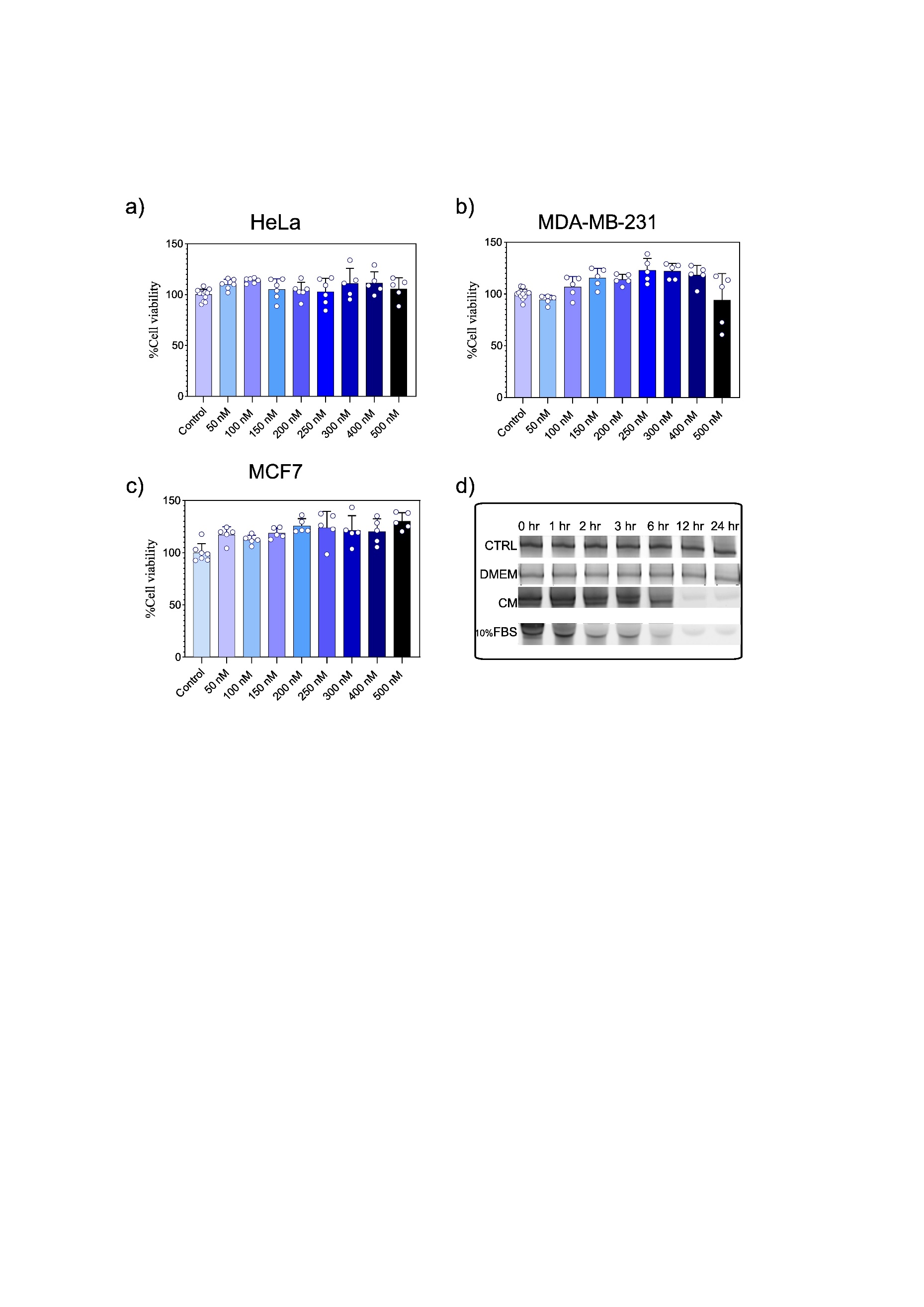

**Supplementary Figure 1**. **Biocompatibility and stability of DNA Td.** (a-c) Cell viability of HeLa (a), MDA-MB-231 (b), and MCF-7 (c) cells treated with increasing concentrations of DNA Td (0-500 nM) for 24 hrs, measured by MTT assay. Data represent mean ± SD. (d) Stability of DNA Td incubated in control buffer, DMEM, Complete DMEM Media (CM), and 10% FBS over 24 h.

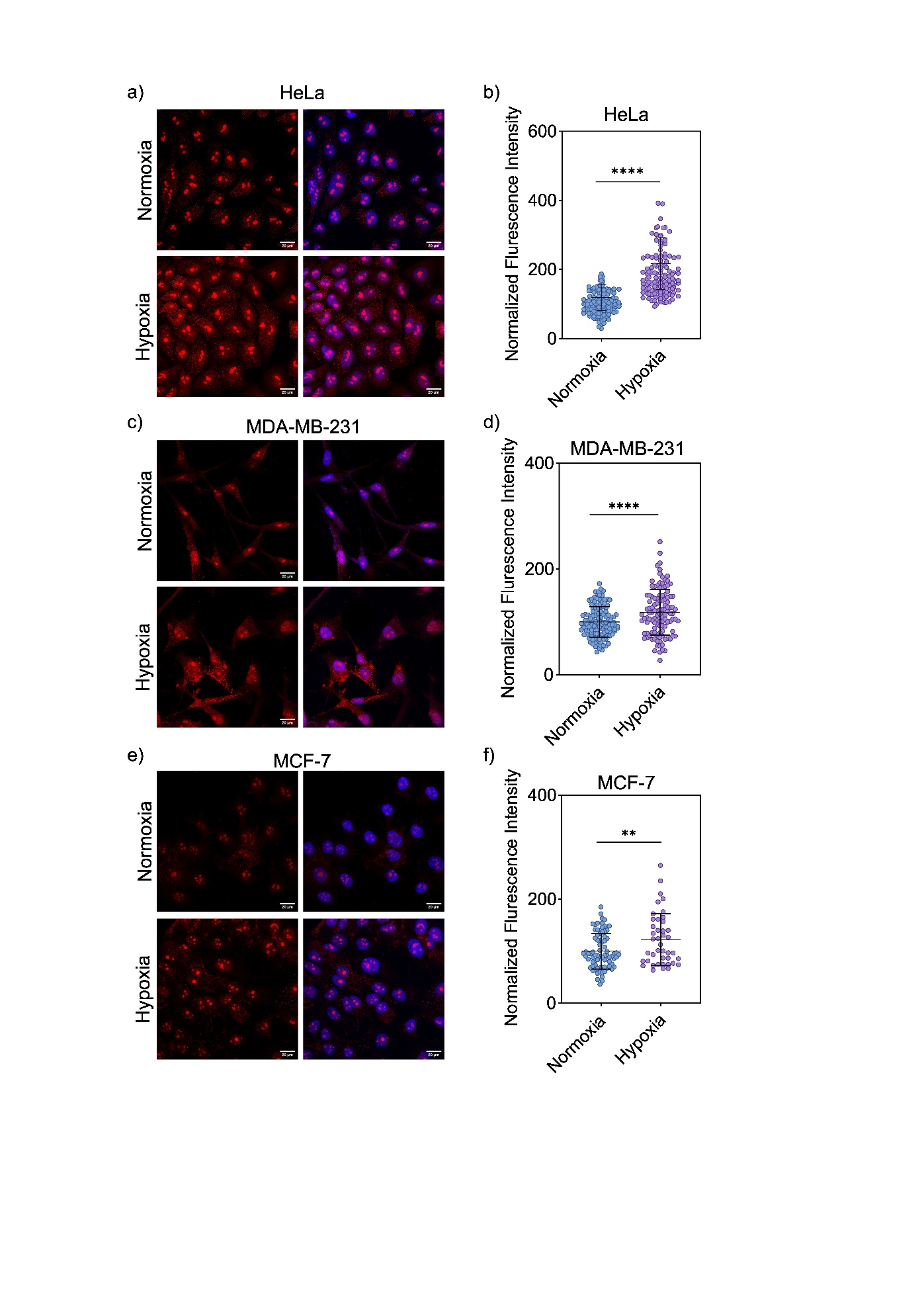

**Supplementary Figure 2. CoCl₂ treatment induces nuclear accumulation of HIF-1α across cancer cell lines.** (a, c, e) Representative immunofluorescence images of HeLa (a), MDA-MB-231 (c), and MCF-7 (e) cells under normoxic and hypoxia-mimetic (100 μM CoCl₂, 24 h) conditions. Left panels show HIF-1α staining (red); right panels show merged images with DAPI-stained nuclei (blue). Scale bars, 20 μm. (b, d, f) Quantification of normalized nuclear HIF-1α fluorescence intensity in HeLa (b), MDA-MB-231 (d), and MCF-7 (f) cells under normoxia versus hypoxia-mimetic conditions. Data represent individual cell measurements (n > 50 cells per condition) with mean ± SD. Statistical significance was determined by unpaired t-test (**P < 0.01, ****P < 0.0001).

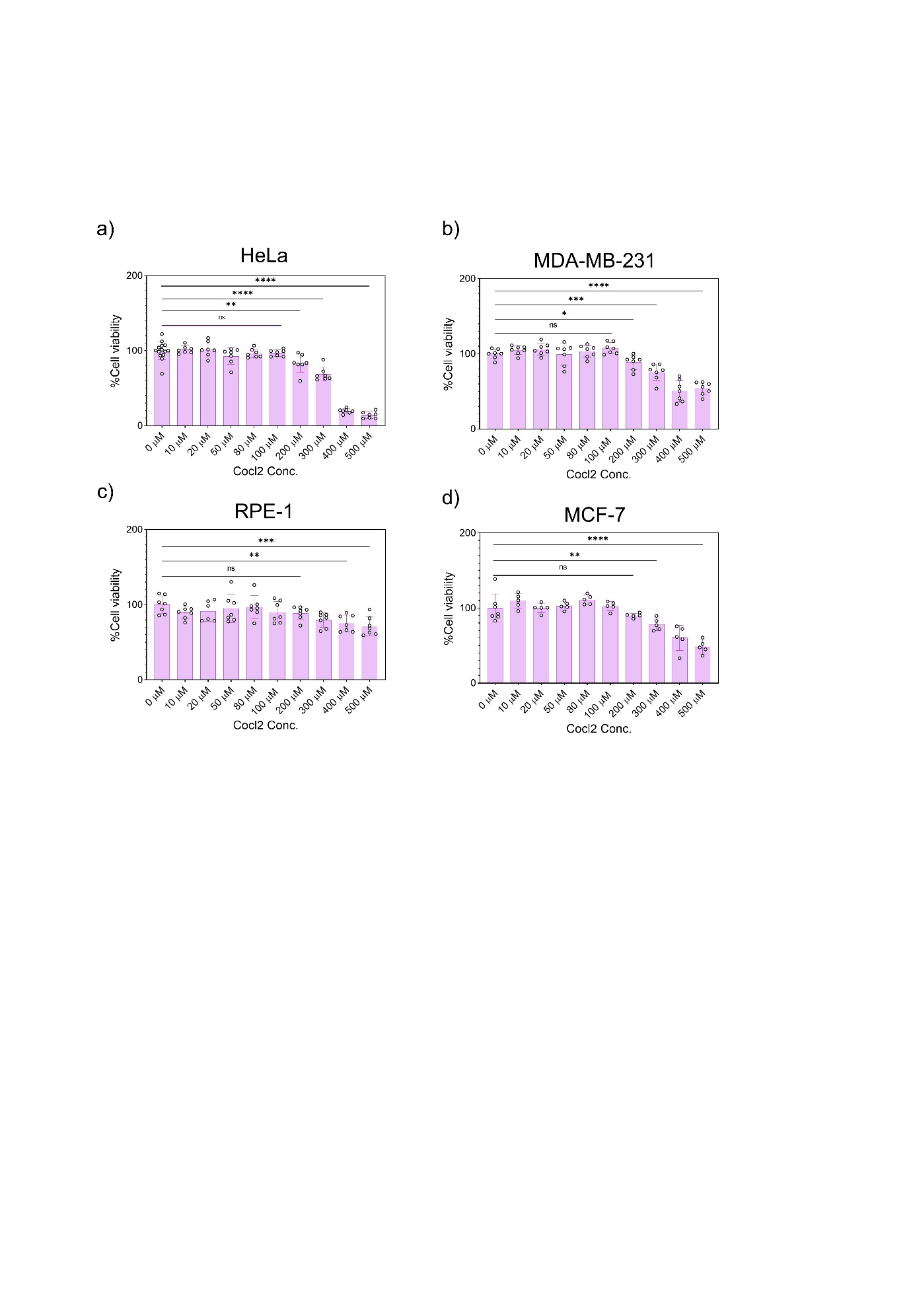

**Supplementary Figure 3. Dose-dependent effect of CoCl₂ on cell viability across cell lines.** (a-d) Cell viability of HeLa (a), MDA-MB-231 (b), RPE-1 (c), and MCF-7 (d) cells treated with increasing concentrations of CoCl₂ (0-500 μM) for 24 h, measured by MTT assay. Viability is expressed as a percentage relative to untreated (0 μM) controls. Data represent mean ± SD. Statistical significance relative to the 0 μM control was determined by one-way ANOVA (ns, not significant; *P < 0.05; **P < 0.01; ***P < 0.001; ****P < 0.0001).

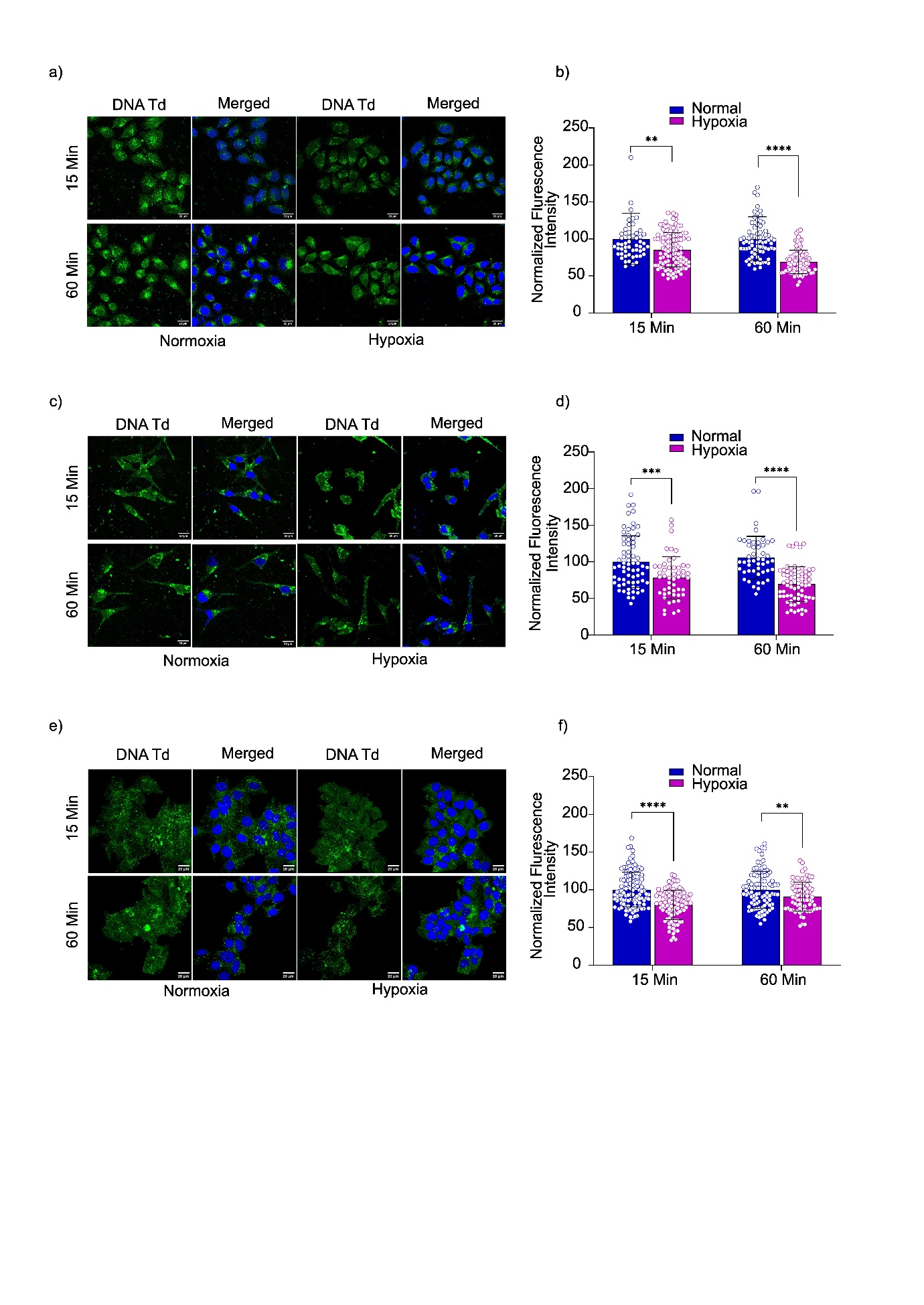

**Supplementary Figure 4: Hypoxia reduces DNA tetrahedron (DNA Td) uptake in different cell lines.** (a, c, e) Representative confocal fluorescence images showing intracellular uptake of fluorescently labelled DNA Td under normoxic and hypoxic conditions at 15 and 60 min. DNA Td fluorescence is shown in green, while nuclei are counterstained with DAPI (blue). (b, d, f) Quantification of normalized intracellular fluorescence intensity at 15 and 60 min under normoxia and hypoxia. Hypoxia significantly reduced DNA Td uptake at both time points across the tested cell lines. Data are presented as individual measurements with mean ± SD. Statistical significance was determined using an appropriate statistical test; **P < 0.01, ***P < 0.001, ****P < 0.0001.

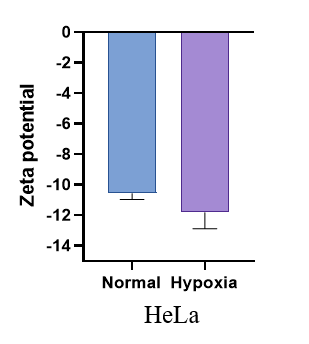

**Supplementary Figure 5:** Zeta potential of the HeLa cells under normoxic and hypoxic conditions.
